# Physiology of PNS axons relies on glycolytic metabolism in myelinating Schwann cells

**DOI:** 10.1101/2020.04.23.049056

**Authors:** Marie Deck, Gerben Van Hameren, Graham Campbell, Nathalie Bernard-Marissal, Jérôme Devaux, Jade Berthelot, Alise Lattard, Jean-Jacques Médard, Benoît Gautier, Patrice Quintana, Juan Manuel Chao de la Barca, Pascal Reynier, Guy Lenaers, Roman Chrast, Nicolas Tricaud

**Affiliations:** INM, INSERM, Université de Montpellier, 80 Rue A. Fliche, 34090 Montpellier, France; Aix-Marseille University, INSERM, MMG, 13385 Marseille, France; Departments of Clinical Neuroscience and Neuroscience, Karolinska Intitutet, Stockholm, Sweden; Département de Biochimie et Génétique, Centre Hospitalier Universitaire, Angers, France; Equipe Mitolab, MITOVASC, CNRS 6015, INSERM U1083, Université d’Angers, Angers, France; I-Stem, UEVE/UPS U861, INSERM, AFM, 91100, Corbeil-Essonnes, France

## Abstract

Despite the lactate shuttle theory, how glial cells support axonal metabolism and function remains unclear. Lactate production is a common occurrence following anaerobic glycolysis in muscles. However, several other cell types, including some stem cells, activatezd macrophages and tumor cells, can produce lactate in presence of oxygen and cellular respiration, using Pyruvate Kinase 2 (PKM2) to divert pyruvate to lactate dehydrogenase. We show here that PKM2 is also upregulated in mature myelinating Schwann cells (mSC) of mouse sciatic nerve. Deletion of this isoform in PLP-expressing cells in mice leads to a deficit of lactate in mSC and in peripheral nerves. This had no detectable consequences on the myelin sheath. However, mutant mice developed a peripheral neuropathy. Peripheral nerve axons of mutant mice failed to maintain lactate homeostasis upon activity, resulting in an impaired production of mitochondrial ATP. Action potential propagation was not altered but axonal mitochondria transport was slowed down, muscle axon terminals retracted and motor neurons displayed cellular stress. Additional reduction of lactate availability through dichloroacetate treatment, which diverts pyruvate to mitochondrial oxidative phosphorylation, further aggravated motor dysfunction in mutant mice. Thus, lactate production through aerobic glycolysis is essential in mSC for the long-term maintenance of peripheral nerve axon physiology and function.

## INTRODUCTION

Energetic metabolism is an essential parameter of neurons’ function and survival (Harris & Attwell, 2012; Attwell & Laughlin, 2001; Alle *et al*, 2009). Indeed, the nervous system consumes a large amount of glucose mainly to allow synapses to function but also organelles and vesicles transport along axons and action potentials firing (Harris & Attwell, 2012). In the peripheral nervous system (PNS), axons grow far from their cell bodies to reach distant targets such as muscles. Maintaining metabolic homeostasis on such a long distance is a real challenge for peripheral nerve axons (Waxman, 1997).

To support neurons in this challenge, myelinating Schwann cells (mSC) cover the large motor axons with a myelin sheath This insulating sheath, resulting from several turns of compacted plasma membrane, allows for ionic and electronic insulation forcing action potentials to “jump” from one node of Ranvier to the next, accelerating the nerve conduction velocity up to 120m/s (Waxman, 1980). Unfortunately, this insulation also hinders the diffusion of extracellular metabolites in the axonal environment mechanically limiting metabolic support to myelinated axons.

However, glial cells have also been shown to provide a direct trophic support to the axons they surround. Indeed, evidence exist for a metabolic coupling between glial cells and neurons in the CNS. Astrocytes and oligodendrocytes play a critical role in this process by metabolizing glucose into lactate and exporting it to the axon as a fuel for axonal mitochondria, a process known as the lactate shuttle (Magistretti & Allaman, 2015; Fünfschilling *et al*, 2012; Lee *et al*, 2012). Recently, mitochondrial respiration was shown to be dispensable for myelinating oligodendrocytes suggesting these cells can use glycolysis for ATP production and produce lactate (Della-Flora Nunes *et al*, 2017; Fünfschilling *et al*, 2012). The disruption of monocarboxylate transporters (MCT), which mediate the traffic of metabolites, induced motor endplate dysfunction (Bouçanova *et al*, 2021; Jha & Morrison, 2020), axonal damages and the degeneration of neurons *in vivo* (Lee *et al*, 2012; Philips *et al*, 2021)..

Lactate production in aerobic conditions has been well studied in cancer cells. Indeed, many type of cancer cells use aerobic glycolysis, also called the Warburg effect, to produce ATP as well as other metabolites required for cell survival and proliferation (Vander Heiden *et al*, 2009). This metabolic shift to the aerobic glycolysis relies on the expression of Pyruvate Kinase M2 (PKM2) isoform instead of PKM1 (Israelsen *et al*, 2013, 2). PKM2 is a master regulator of glycolysis cumulating metabolic and non-metabolic functions as a protein kinase and transcriptional coactivator (He *et al*, 2017, 2). How PKM2 expression leads to more lactate is not clear. However, as PKM2 slows down pyruvate synthesis compared to PKM1, this may allow lactate dehydrogenase to metabolize pyruvate into lactate (Hanahan & Weinberg, 2011; He *et al*, 2017; Frezza & Gottlieb, 2009).

Recently, cases of reversible peripheral neuropathy have been observed following treatment with dichloroacetate (DCA), a chemical compound acting on lactate level in cells (James & Stacpoole, 2016; Tataranni & Piccoli, 2019). Indeed, DCA is indirectly activating pyruvate dehydrogenase (PDH) and increasing the mitochondrial uptake of pyruvate, depleting cellular lactate. While the use of this compound to prevents cancer cells growth in several tumors is controversial (Stacpoole, 1989; Tataranni & Piccoli, 2019), it is also indicated to treat acute and chronic lactic acidosis and diabetes (James *et al*, 2017). DCA side effects suggest the importance of lactate balance in the PNS function.

Here, we investigated the role of lactate in the physiology and function of axons and mSC of the PNS through the deletion of PKM2 in mSC (mSC–PKM2). Our results revealed the delicate equilibrium of lactate homeostasis in myelinated fibres of peripheral nerves and the critical trophic role of mSC in the support of axonal function in PNS.

## RESULTS

### PKM2 expression promotes aerobic glycolysis in mSC

Firstly, to investigate the metabolic status of SC, we examined the expression and localization of PKM1 and PKM2 isoforms in mouse sciatic nerves using RT-_q_PCR and immunohistochemistry. PKM1 mRNA was downregulated during sciatic nerve maturation from postnatal day (P) 2 to P28 while PKM2 expression increased at the same time (**Fig. 1A**). At non mature ages of SC, P4 and P15, PKM1 and PKM2 were both expressed in mSC (characterized by E-cadherin staining)(Tricaud *et al*, 2005) that surrounded axons (characterized by 2H3 staining, **Fig. 1B a,b,c,d**). When SC were mature and formed myelin, at P30 and 5 months, PKM2 replaced PKM1 in mSC (**Fig. 1B e,f,g,h**), in particular in the perinuclear region (**Fig. S1**), suggesting mSC enters the aerobic glycolysis metabolic mode when the myelinated fibers are mature.

**FIGURE 1.**
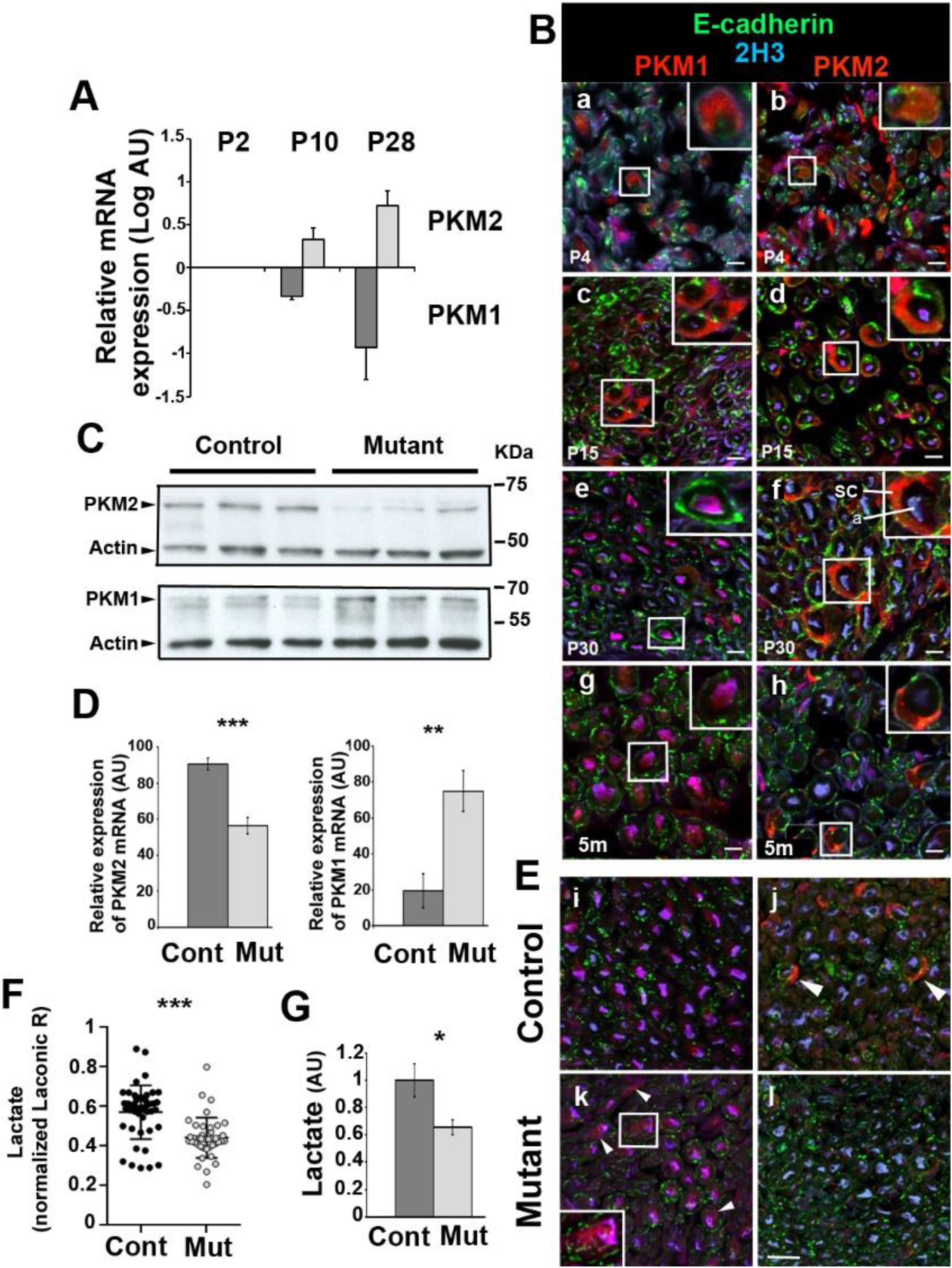
PKM2 is efficiently deleted in mature mSC of mutant mice. **A**-Quantitative RT-PCR on sciatic nerve extracts shows decreased PKM1 and increased PKM2 expressions during mSC maturation. Normalized on P2 values. n=4 animals. **B**-Immunostaining on mouse sciatic nerve sections shows decreased PKM1 (left panels) and increased PKM2 (right panels) expressions in mSC. m: months. Sc: Schwann cell; a: axon. **C**-Western blots on sciatic nerve extracts show lower PKM2 and higher PKM1 amounts in Mutant versus Control mice sciatic nerves (2 months old). Actin serves as loading control. n=3 mice. **D**-Quantitative RT-PCR on sciatic nerve extracts shows decreased PKM2 and increased PKM1 expressions in Mutant (Mut, n=7) versus Control (Cont, n=5) mouse sciatic nerves (samples from 3 and 12 months old mice were pooled). **E**-Immunostainings on sciatic nerve sections show PKM2 lower in Mutant mSC (panel l) while PKM1 is higher (arrowheads and Insert panel k)(2 months old). **F**-The ratio of Laconic probe was measured in mSC of Mutant (n=11 mice, 41 cells) and Control (n=11 mice, 45 cells) anesthetized mice and normalized to the mean of Control values (7 to 10 weeks old mice). **G**-Biochemical measure of lactate shows a reduced concentration in Mutant versus Control sciatic nerves (n=6 mice)(samples from 4 and 12 months old mice were pooled). All scale bars = 10μm. Two-tailed Student t-test. Error bars represent SD (A, D) or SEM (F). AU: arbitrary unit.

### PKM2 deletion in mSC leads to a reorganization of the metabolism in the nerve

The detected expression of PKM2 in mature mSC suggested that they were producing lactate, which raised the possibility that they provided a trophic support to axons. To test this hypothesis, we first performed a time-restricted conditional deletion of PKM2 isoform in myelinating glia of mice. A mouse strain expressing floxed PKM2-specific exon 10 alleles (Israelsen *et al*, 2013) was crossed with a strain expressing the Cre recombinase under the inducible and myelinating cell-specific promoter PLP1-ERt (Doerflinger *et al*, 2003). Injecting tamoxifen in Cre positive/*PKM2*^fl/fl^ (mutant) and Cre-negative/*PKM2*^fl/fl^ (control) littermates at 1 month of age resulted in a significant decrease of PKM2 expression in sciatic nerves of mutant as detected by immunoblotting, quantitative RT-PCR and immunostaining (**Fig. 1C,D,E**). The same experiments indicated that PKM1 was re-expressed in PKM2-deleted mSC (**Fig. 1D,E**), suggesting a compensatory mechanism to maintain energy production.

Next, we expressed a lactate-detecting fluorescent probe (Laconic, **Fig. S2**, San Martín et al., 2013) in mSC of the mouse sciatic nerve *in vivo* using an AAV9 vector (Gonzalez *et al*, 2014). Mutant and control mice expressing the probe were then anesthetized and their nerves imaged using a multi-photon microscope to measure the relative amount of lactate in mSC of living mice. Cells of mutant mice showed significantly less lactate than mSC of control mice (**Fig. 1F**). In addition, biochemical measurements showed that this deficit of lactate in mSC resulted in a global reduction of the amount of lactate in mutant mice nerves compared to control ones (**Fig. 1G**).

To analyze in a broader way the influence of PKM2 deletion in mSC on the metabolic status of the nerve, we performed a targeted metabolomic screen on mutant and control mouse nerves. We observed a decrease of some acylcarnitines in mutant mice (**Fig. S3**), without modification of the acylcarnitine/carnitine ratio (0.2707 vs 0.2739 in control and mutant respectively, P=0.77 two –tailed Student T-test). These acylcarnitines are involved in the transport of fatty acids across mitochondrial membranes before their degradation in carnitine and acetyl-CoA (Indiveri *et al*, 2011). Therefore, the maintenance of the ratio coupled to the decrease of acylcarnitines indicated a higher turnover, suggesting an increase of the mitochondrial activity. Moreover, as mitochondrial dysfunction following deletion of TFAM1 in mSC of mice has been shown to dramatically increase acylcarnitines (Viader *et al*, 2013), our data suggested the opposite effect in mutant mice, i.e. the upregulation of mitochondrial activity.

### Lactate homeostasis and ATP production are impaired in axons of PKM2-SCKO mice

According to the lactate shuttle theory this decrease of lactate should also result in a shortage of axonal lactate. We investigated this assumption by expressing Laconic in peripheral axons using an AAV9 vector injected intrathecally in young pups (**Fig. S4A**). After tamoxifen induced recombination we performed live-imaging of probe-labeled axons that cross the saphenous nerve in anesthetized mice (**Fig. S4A**). While in resting conditions no difference could be seen between genotypes (**Fig. S5A**), when nerves were challenged with electrical stimulations to generate action potentials in type A fibers, axonal lactate increased shortly after the stimulation in control but dropped in mutant (**Fig. 2A**). In the long term, control mice axons were able to maintain their lactate homeostasis while mutant mice axons could not (**Fig. 2A**, red line).

**FIGURE 2.**
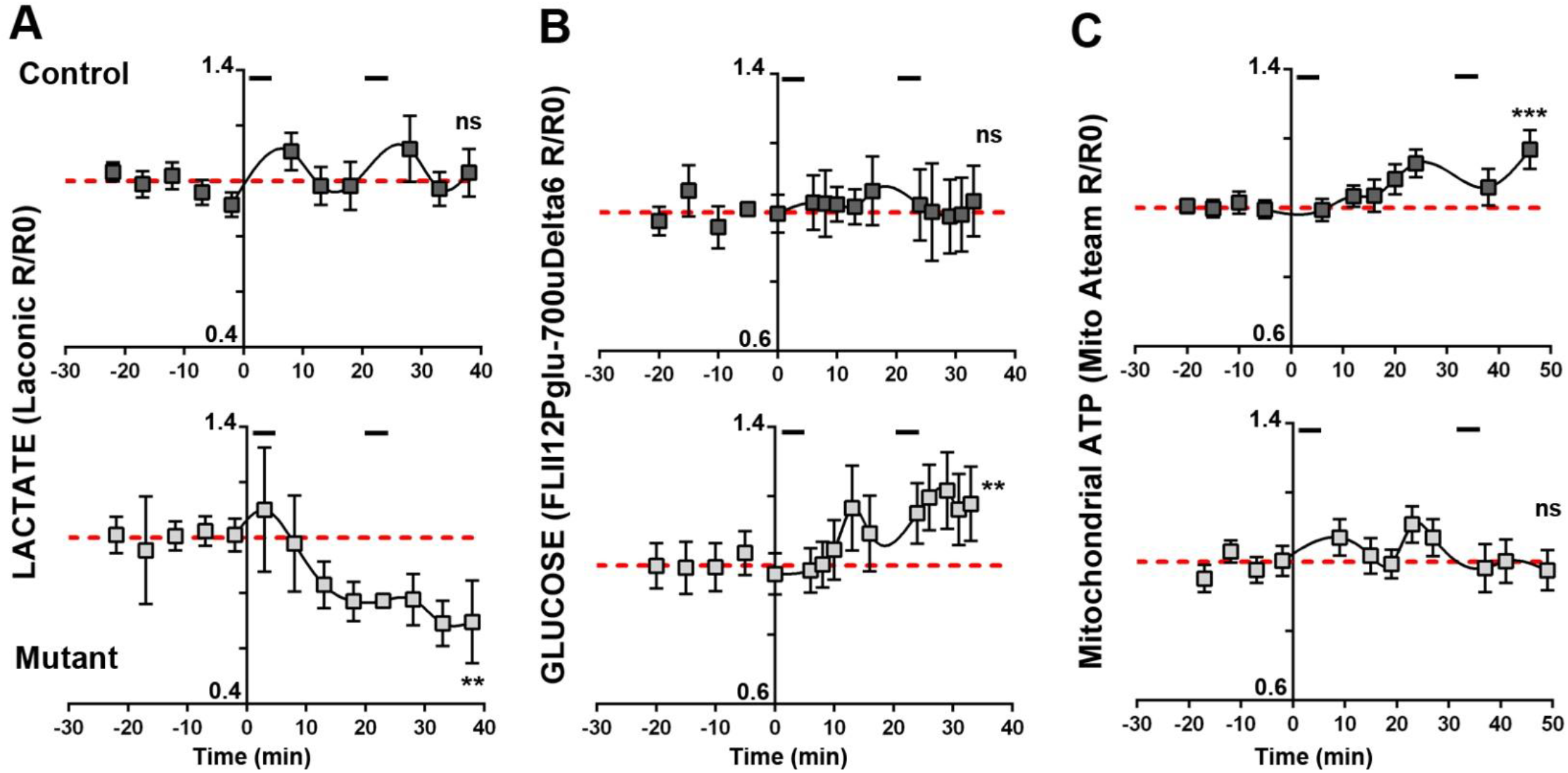
Mutant mice axons display an altered metabolic response to electrical stimulations. The ratio of fluorescent probes (R) was normalized over the average ratio at time points before the first stimulation (R0)(basal value). Segments at the top of each Control mice graphs show stimulation periods. Lines show the basal value line. Statistical analysis shows the test for linear trend following one-way ANOVA analysis. Error bars represent SEM. **A-** n= 20 axons, 7 mice for Control and 7 axons, 4 mice for Mutant. **B-** n= 6 axons, 3 mice for Control and 9 axons, 3 mice for Mutant. **C-** n= 16 axons, 5 mice for Control and 14 axons, 5 mice for Mutant.

Beside lactate, another way axons may feed their mitochondria is glucose-derived pyruvate. Thus, we used a glucose-specific probe (FLII12Pglu-700uδ6, **Fig. S4B**) (Takanaga *et al*, 2008) to investigate glucose level in axons in the same conditions. While in resting axons a similar amount of glucose was found in both genotypes (**Fig. S5B)**, upon electrical stimulations glucose increased in axons of mutant mice while it remained stable in control mice (**Fig. 2B**), suggesting that axons of mutant mice mobilized glucose instead of lactate upon stimulations. We finally investigated the production of ATP by axonal mitochondria *in vivo* using a mitochondria-targeted fluorescent probe detecting ATP (ATeam, **Fig. S4C**) (Tsuyama *et al*, 2013). Again, no difference could be seen in resting conditions (**Fig. S5C)**. However, when axons were stimulated mutant mice mitochondria failed to increase ATP production while this production increased in control mice (**Fig. 2C**). Thus, in absence of aerobic glycolysis in mSC, lactate homeostasis and mitochondria ATP production are impaired in electrically active axons, despite the mobilization of glucose.

### Behavior impairment and neuromuscular junction loss in PKM2-SCKO mice are not due to demyelination

Since we observed a deficit in energy production in axons, we tested whether this had an impact on motor capacities of mutant mice. A longitudinal analysis revealed a significant deficit in both Rotarod and grip tests in mutant mice (**Fig. 3A,B**), suggesting a direct influence of the metabolic changes observed in mutant mice axons on their motor abilities. We also measured the nerve conduction velocity of sciatic nerves, and noticed it was not altered in mutant mice **(Fig. 3C**) indicating that the myelin sheath was not affected. Accordingly, electron microscopy analyses showed correctly myelinated axons and a unvarying g-ratio in mutant mice (**Fig. S6**). The conduction of action potentials along myelinated axons was also not altered as the electrophysiological properties of sciatic nerve axons *ex vivo* were maintained even at very high firing frequencies (**Fig. S7, Table S**). Motor neuron number did not change in the spinal cord (**Fig. 3D**) but they displayed a higher expression of cleaved Caspase 3 (**Fig. 3E and S8**), a marker of neuronal stress and axonal degeneration (Mukherjee & Williams, 2017). Indeed, tracking axonal mitochondria *in vivo* revealed a decrease of their movements in mutant mice both before and after electrical stimulation (**Fig. 3F**). In addition, mutant mice showed significantly more denervated neuromuscular junctions with intact postsynaptic structures (**Fig. 3G,H**) in the gastrocnemius muscle, suggesting the retraction of axon terminals. Taken together, these data indicated that PKM2 deletion did not affect the maintenance of the myelin sheet but resulted in a motor distal neuropathy in mice. This indicated that aerobic glycolysis is required in mSC for the maintenance of axonal trafficking and neuromuscular junctions.

**FIGURE 3.**
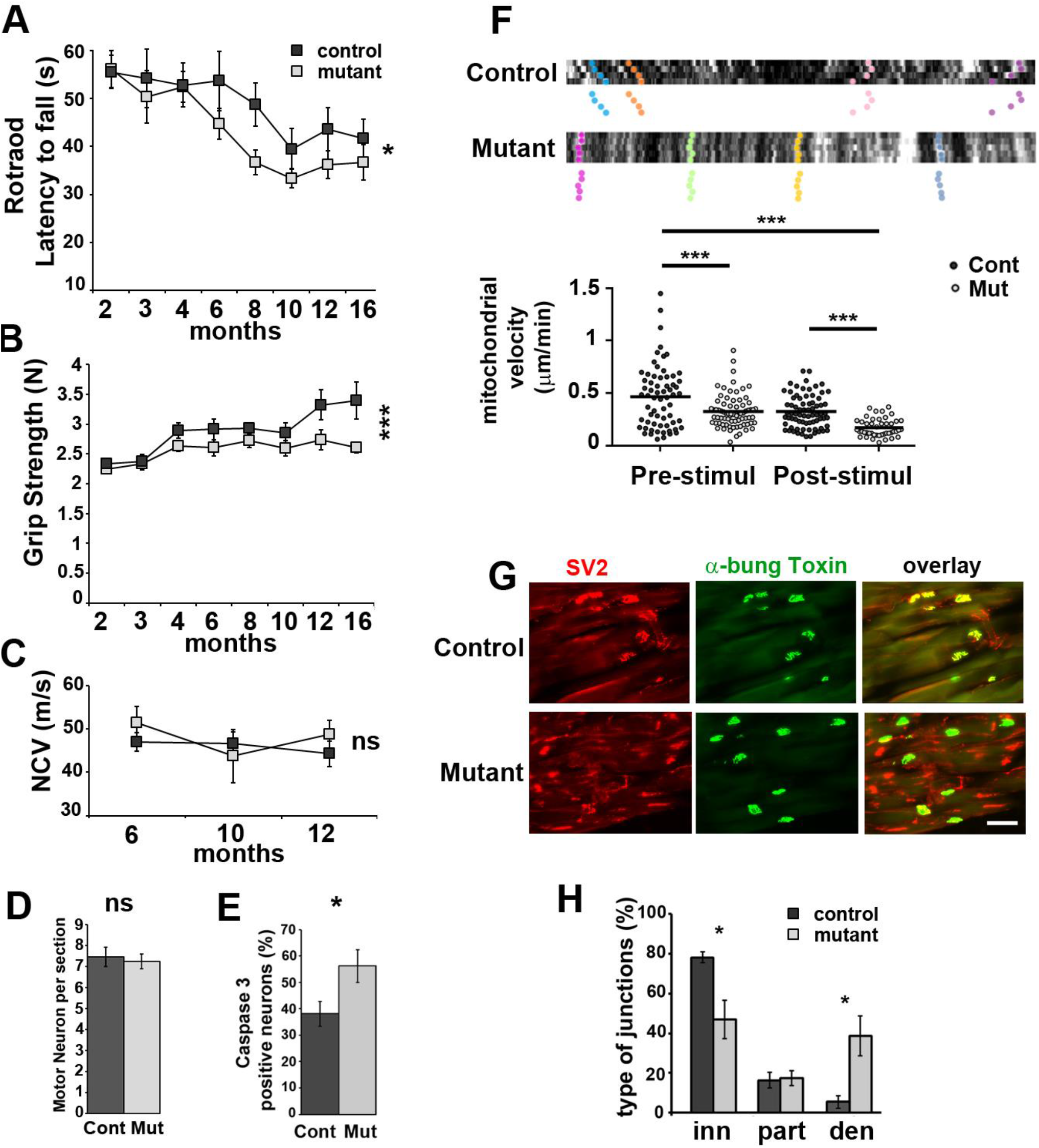
Mutant mice show motor deficit and display axonal defects. **A-** Rotarod latency to fall and **B**-grip strength were measured between 2 and 16 months postnatal on Control (n=21) and Mutant (n=25) mice. Two-way non repeated measures ANOVA Sidak’s post-hoc tests. **C-** Nerve conduction velocity (NCV) measured in Control and Mutant mice (n=10 each). Two-way non repeated measures ANOVA Sidak’s post-hoc tests. **D**-Spinal cord sections of Mutant (Mut) and Control (Cont) mice (12 months) were stained with cresyl violet and motor neurons were counted (see Fig. S8). n= 3 animals (14 to 30 sections). Two tailed Student t-test. **E-** Spinal cord sections of Mutant (Mut, n=19 images, 3 animals) and Control (Cont, n= 12 images, 2 animals) mice were immunostained for Caspase 3, ChAT and Neurofilament (Fig. S8) (12 months old). Caspase 3 positive neurons in percentage of Neurofilament positive neurons. Two tailed Student t-test. **F-** Axonal mitochondria labelled with mito-Dsred2 were imaged *in vivo* before and after electrical stimulation. **Upper panels** show typical kymographs. Tracked mitochondria are shown with colour dots. Mutant mice mitochondria follow a straight pattern indicating they are immobile or slowly moving. **Lower panel**: Mitochondria speed was plotted according to the genotype before and after stimulation. One-way ANOVA Tukey’s post-hoc test. n are provided in Material and Methods. **G-** Gastrocnemius muscle neuromuscular junctions of 12 months old Mutant and Control mice were stained for presynaptic SV2 and postsynaptic acetylcholine receptor with FITC-α-bungarotoxin (α-bung). Scale bar= 100μm. **H-** Innervated (inn, complete overlap), partially innervated (part, partial overlap) and denervated (den, no overlap) junctions were counted on sections as shown in D. Two tailed Student t-test. n= 4 mice (12 months). Error bars represent SEM.

### Long-term maintenance of axons function relies on lactate feeding by SC

To investigate further the role of lactate intake in axons’ long term maintenance, we treated mutant and control mice with dichloroacetate (DCA), a drug that promotes mitochondrial consumption of pyruvate and decreases the availability of lactate (**Fig. 4A**) (Stacpoole, 1989). DCA treatment has been considered for patients suffering from congenital lactic acidosis and for cancer treatment but it can lead to a reversible peripheral neuropathy primarily affecting axons in humans and rodents (James & Stacpoole, 2016). Over the seven weeks of treatment, Rotarod performance decreased in treated mice independently of the genotype (**Fig. 4B**) with no effect on the myelin sheath as the nerve conduction velocity did not significantly change (**Fig. 4C**). However, when the treatment stopped, control mice recovered over 3 weeks on the Rotarod while mutant mice did not (**Fig. 4B**). This indicated that axonal function was definitely damaged in these mice, confirming that aerobic glycolysis is required in mSC for the long-term maintenance of peripheral axons’ physiology.

**FIGURE 4.**
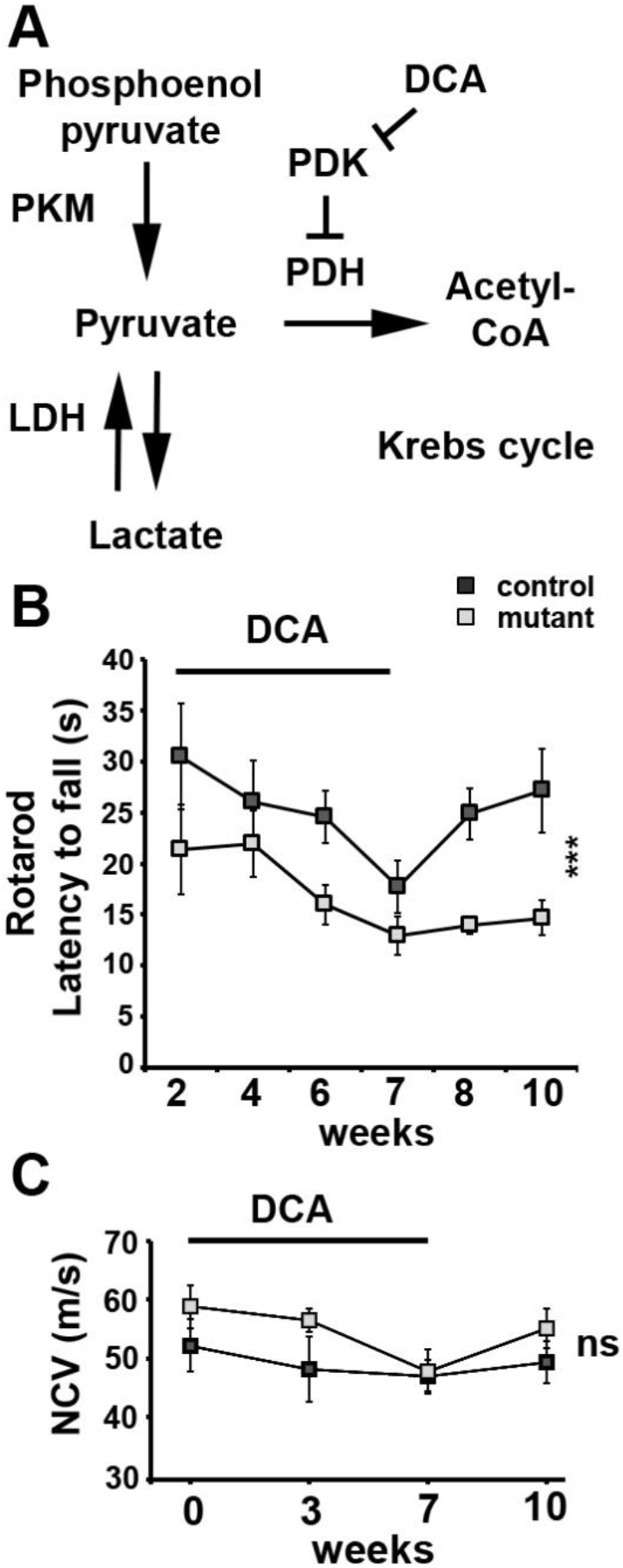
PKM2 deletion in mSC impairs motor performance recovery after DCA treatment. **A-** Metabolic pathways directly affected by the manipulations. LDH: lactate dehydrogenase; PDH: pyruvate dehydrogenase; PDK: PDH kinase. **B-** Rotarod latency to fall was measured in 9 months old Control (n=5) and Mutant mice (n=5) during a 7 weeks daily treatment with DCA and then during a three weeks recovery period without treatment. Two-way non repeated measures ANOVA Sidak’s post-hoc test from week 7 to week 10. **C**-Nerve conduction velocity (NCV) measured in 9 months old Control (n=5) and Mutant mice (n=7) at 0, 3 and 7 weeks of DCA treatment and 3 weeks after stopping the treatment (week 10). Two-way non repeated measures ANOVA statistical tests, Sidak’s post-hoc tests over the 4 time points. Error bars represent SEM.

## DISCUSSION

While the role of the lactate in the metabolic crosstalk between axons and the surrounding glia has been recognized for a long time in the CNS (Harris & Attwell, 2012), its role remains unclear in the PNS. The presence of lactate in peripheral nerves had been detected and *ex vivo* experiments had shown that it could support action potentials propagation along axons (Brown *et al*, 2012). However, the cellular origin of the released lactate and its relevance as an energy substrate for axons remains largely unclear (Stassart *et al*, 2018). By deleting PKM2 in mSC *in vivo* we showed that part of this nerve lactate is provided by mSC using aerobic glycolysis and this lactate is required for axons to maintain their metabolic homeostasis and functions over time.

Unexpectedly, and to the opposite of cancer cells, preventing mSC performing aerobic glycolysis had very little impact on its own physiology and biology. Indeed, no defect could be detected in the myelin sheath of PKM2 mutant mice and the nerve conduction was not affected. A slight but significant shift occurred in the amount of some acylcarnitines without any alteration of the carnitine/acylcarnitine ratio, suggesting a higher turnover of these compounds that shuttle lipids into mitochondria for β-oxidation. According to the literature we interpreted these data as the sign of a higher metabolic activity of mSC mitochondria. This is consistent with the upregulation of PKM1 expression we observed in mutant mice nerves as the higher enzymatic activity of this isoform promotes pyruvate production and its use in the mitochondrial citric acid cycle and respiration. In the absence of PKM2, mSC metabolism probably shifted to more oxidative phosphorylation. As mSC cannot be dispensed of mitochondrial respiration (Della-Flora Nunes *et al*, 2017; Fünfschilling *et al*, 2012), this shift is likely to be mild for the cell which is still able to produce myelin and maintain it. However, as evidenced by our results the shutdown of aerobic glycolysis in mSC is clearly deleterious for axonal maintenance. Therefore, we conclude that mSC use simultaneously both mitochondrial respiration for myelin production and maintenance and aerobic glycolysis to support axons survival.

The challenge in the analysis of metabolic crosstalk in a tissue is to distinguish between the metabolisms of the different cell types. In addition, nerve injury or section immediately modifies both axonal and glial mitochondria physiology (Kerschensteiner *et al*, 2005). To overcome these challenges, we chose to use the genetically encoded fluorescent probes delivered specifically to axons or mSC *in vivo* to follow a few critical metabolites in real time in physiological conditions. These probes such as lactate sensor *Laconic*, the glucose sensor *FLII12Pglu-700uδ6* and the ATP sensor *ATeam* have been extensively characterized in several *in vitro* and *in vivo* models and we confirmed that variations could be observed in the nerve *in vivo*. Although this gave us a limited view of all the metabolic changes that may have occurred in mutant cells, we could nevertheless draw some decisive conclusions.

Firstly, we observed that the metabolic homeostasis is really efficient in peripheral nerves. While electrically activated axons show radical alterations in mutant mice in just a few dozens of minutes in particular for lactate, no change could be detected in resting axons of anesthetized animals disregarding the genotype. In this regard, axonal homeostasis is much more efficient than mSC homeostasis as lactate levels remained significantly lower in these cells in mutant mice than in control mice. However, axonal homeostasis was not sufficient to buffer the metabolic changes that occur following physiological stimulations. Indeed, while axonal lactate remained apparently unaltered after stimulations in control mice, it sharply and steadily dropped in mutant mice axons. This revealed that actually the maintenance of lactate levels in control mice axons is due to a significant uptake of mSC lactate to compensate for a severe consumption during axonal activity (Rich & Brown, 2018). Therefore, lactate produced by mSC is a critical fuel for mitochondria of actively-firing myelinated axons. Indeed, even the mobilization of glucose in stimulated axons was not sufficient to sustain an adequate ATP production in mitochondria. The involvement of the mSC lactate was definitively confirmed by the DCA treatment. Mutant mice were unable to recover their performances on the Rotarod following the draining of lactate supplies, while mice that could produce lactate in their mSC through aerobic glycolysis recovered. This underlines the dependence of myelinated axons to mSC lactate as all the other sources of energy substrates such as glucose, glycogen or glutamine are not directly altered by DCA. The reason for this dependence is puzzling but one possibility could be the swift accessibility of this glial lactate pool and its readiness for mitochondrial respiration while using glucose may require more time and enzymatic resources in the axons to generate pyruvate. This concept is supported by the recent discovery of a transient upregulation of glycolysis in mSC and the involvement of MCT to promote axons survival in an peripheral nerve injury model (Babetto *et al*, 2020).

The main macroscopic phenotype resulting from PKM2 deletion in mSC is a motor weakness observed through Rotarod and grip test in the absence of demyelination. This was not significantly detectable before 6 months suggesting a late onset of the distal motor neuropathy. Taken together, this is characteristic of axonal CMT diseases in mice. Moreover, this neuropathy did not result from motor neurons death but from an axonal dysfunction illustrated by a decreased mitochondrial motility and retracted neuromuscular terminals. This retraction is likely to be the main cause of the motor weakness. Mitochondrial migration, which is essential for the maintenance of the synapses, in particular in motor neurons (Marinković *et al*, 2012), relies on the ATP produced in mitochondria. In a previous work we showed that shutting down ATP synthase activity steadily slowed down mitochondrial movements in mSC *in vivo* (Gonzalez *et al*, 2015). Taken together, our results suggest that, the failure of mutant mice axonal mitochondria to increase their ATP production following physiological activity is the likely initial cause of the axonal dysfunction.

However, axonal firing properties were not significantly altered by this mitochondrial failure. This was unexpected because a largely accepted idea is that axonal ATP, and therefore axonal mitochondria, are required to maintain the membrane negative potential that allows depolarization. Actually, more recent analysis of neuronal energetics indicates that the energy cost of maintaining axon membrane potential is not that important, especially in myelinated fibers (Harris & Attwell, 2012). Therefore, the production of ATP by mutant mice axonal mitochondria may be sufficient to support this cost. Nevertheless, our data clearly indicate that mitochondrial ATP production and adaptation to the axon activity is critical for axonal maintenance.

Matching the energy demand of axons is critical to ensure the long-term maintenance of the nervous system. Indeed, increasing evidences indicate that an unbalanced energy supply to axons is a capital factor of neurodegenerative diseases such as ALS (Vandoorne *et al*, 2018), Parkinson and Alzheimer diseases (Camandola & Mattson, 2017) or leprosis *Mycobacterium Leprae*-induced peripheral neuropathy (Medeiros *et al*, 2016). However, the molecular basis of this support remains unclear. The present data show that axonal dysfunction may be a direct result of perturbed metabolic support by glial cells such as mSC. In this regard, the dramatic effect of

DCA treatment on control and mutant mice motor performances is striking and the inability of mutant mice to recover from this treatment is challenging. Indeed, DCA is proposed as a long-term treatment for several cancers and other diseases such as acute and chronic lactic acidosis and diabetes. Our data suggest that drugs targeting aerobic glycolysis metabolism as a treatment for these diseases may have a singular and deleterious effect on the nervous system and in particular on the PNS. Therefore, further and finer characterization of the axon/glia metabolic crosstalk is required in particular in neurodegenerative diseases. In this regard, the cancer-like nature of this glial metabolism constitutes a risk for the nervous system regarding the deleterious effects of anticancer drugs that affect cell metabolism.

## Supporting information

Raw data and statistical analysis

## Acknowledgments

We thank H.Boukhaddaoui, C.Sar, V.Baecker and the imaging facility MRI, the INM animal facility, L.Diouloufet and C. Cazevieille. European Research Council grant (FP7-IDEAS-ERC #311610), INSERM AVENIR, EpiGenMed Labex, The Neuromuscular Research Association Basel, Swedish StratNeuro program, Swedish Research Council grant (#2015-02394), and AFM-Téléthon (#20044).

## Author contributions

Conceptualization: NT; Formal analysis: MD, GvH, JJM, JMCB; Funding acquisition: RC, NT; Investigation: MD, GvH, NBM, JD, GC, JB, AL, JJM, BG; Methodology: PQ; Project administration: PR, GL, RC, NT; Supervision: NT; Validation: MD, GvH, PR; Visualization: MD, GvH, JJM, NT; Writing – original draft: MD, GvH, NT; Writing – review & editing: all co-authors.

## List of Supplementary Materials

Materials and Methods Fig S1 – S8

Table S

