## Supplementary data for "Physiology of PNS axons relies on glycolytic metabolism in myelinating Schwann cells"

**Supplementary Materials:**

#### Material and Methods:

##### Animals

Schwann cell specific ablation of PKM2 in adult mice (*Plp1-cre$^{ERT}$;PKM2$^{f/f}$*) was obtained by crossing *Plp1-cre$^{ERT}$* (JAX: 005975)(*12*) mice with *PKM2$^{f/f}$* (JAX: 024048) (*11*) to generate *Plp1-Cre$^{ERT}$;Pkm2$^{+/f}$* animals, which were in turn backcrossed with *PKM2$^{f/f}$* mice. All transgenic lines were kept in C57/Bl6 background. Genotyping for all mutants was performed by PCR strategies using standard procedures and appropriate primers from Jackson Laboratories. Animals were kept under French and EU regulations, following recommendations of the local ethics committee.

*Tamoxifen administration:* The recombination of the floxed allele was induced by intraperitoneal injection of tamoxifen (Sigma, T-5648 ) dissolved in corn oil (Sigma, C-8267) once every 24 hours for a total of 5 consecutive days, in 1 month animals at 180 µg/ gram mouse weight.

*DCA administration:* dichloroacetate (DCA, Sigma) dissolve in water was administrated by gavaging six days per week during seven weeks, at 500 mg/ kg mouse weight.

##### Behavioural studies

*Rotarod tests:* Locomotor coordination was performed with a rotating rod apparatus (Bioseb). Mice from 2 to 12 month were placed on the rotating rod and challenged using the following step: on day 1, mice was familiarized to the behavioral room and learned to stay on the Rotarod at constant speed of 8 rpm for 2 min. At day 2, locomotion performance was assessed on the accelerating Rotarod with a rate of 4 to 40 rpm for 2 min. Latency to fall was recorded on 3 trials separating by at least 15 min pauses. Data are expressed as the means from the 3 trials for each animal, normalized according to animal weight, +- standard error of the mean (SEM).

*Griptest:* muscular strength for the 4 limbs was assessed with a grip test apparatus (Bioseb). Mice were held by the tail and allowed to grab the grid and then pulled backwards in a horizontal plane. The maximum force (measured in Newtons) applied to the grid was recorded. Data are expressed as the means from 3 trials for each animal, normalized according to animal weight, +- standard error of the mean (SEM).

*Motor nerve conduction velocity measurement:* mice were anesthetized with 2% isoflurane and maintain at 37°c on a hotplate. Left and right sciatic nerves were successively stimulated at the sciatic notch (proximal stimulation) and the ankle (distal stimulation) via a pair of steel needle electrodes (AD Instruments, MLA1302) with supramaximal pulses (7V) of 0.05 milliseconds delivered using a PowerLab 26T (AD Instruments ML4856). The distance between the 2 sites of stimulation was measured alongside the skin surface with fully extended legs. The latencies of the CMAP were recorded with a second pair of electrodes inserted between the digits of the hind paw and measured from the stimulus artefact to the onset of the M-waves deflection. NCV was calculated by dividing the distance between sciatic notch and ankle sites of stimulation by the subtraction of the distal latency from the proximal latency.

##### Immunohistochemistry

For immunohistochemistry on cryosection, nerves were dissected and fixed 2hours in PFA4% at 4°C. For spinal cord and DRG, mice were first perfused transcardially with PBS, and spinal cord and DRG were dissected and fixed in PFA4% overnight or 10min respectively. All tissues were then transferred to 30% sucrose overnight before embedding into Optimum Cutting Temperature

(OCT, Tissue-Tek) medium. Cryostat sections of nerves or spinal cord (12 µm) were dried at room temperature (RT) for 15 min, washed in PBS, incubated 1h at RT in a blocking solution (0.25% Triton X-100 and 10% normal Goat serum in PBS) and incubated with primary antibodies in blocking solution overnight at 4°C. For immunochemistry on teased fibers, nerves were isolated from mice, fixed for 10 min in Zamboni's fixative, washed in PBS and subsequently teased on glass slides. Slides were then dried overnight at RT and immunostained as described above. Primary antibodies: PKM1: 1/2500 (NBP2-14833, Novus Biologicals); PKM2 1/500 (SAB4200095, Sigma Aldrich); 2H3:1/80 (DSHB); Cleaved caspase3 1/400 (Cell Signaling); Neurofilament SMI32 1/1000; E-Cadherin 1/300 (MABT26, Merck); SV2 1/100 (DSHB). Secondary donkey antibodies coupled to Alexa Fluor 488, Alexa Fluor 594, or Alexa Fluor 647 1/1000 (Invitrogen) and DAPI 1/1000 (Molecular Probes). Imaged were acquired with a Zeiss confocal microscope LSM710.

For neuromuscular junctions immunostaining, gastrocnemius muscles were dissected and fixed 20 minutes in PFA4% and incubated in 25% sucrose solution at 4°C for 24 h. Tissues were embedded in OCT and stored at -80°C before processing. Neuromuscular Junctions (NMJ) occupancy was quantified on 25-µm thick longitudinal gastrocnemius sections. Muscle sections were incubated overnight at 4°C in blocking solution (2% BSA, 10% NGS, 0.1% Triton and PBS) with mouse anti-SV2 (Developmental Hybridoma Bank, 1/50). Sections were next incubated with goat anti-mouse IgG1 Cy3-conjugated antibody (Jackson ImmunoResearch, 1/500) and Bungarotoxin-488 (Molecular Probes, 1/500), and mounted in Mowiol mounting medium. NMJ were imaged on Axioplan fluorescence microscope (Zeiss) with a 20x objective.

Images that did show at least 5 Neurofilament positive cells were not included in the statistical analysis.

**Histology staining**

Cresyl violet staining was undertaken on PFA perfused spinal cord. Cresyl violet solution was prepared with 0.1g cresyl violet acetate in 100ml distilled water and 250µl glacial acetic acid and filtered before use. Slides were kept at -20˚C and removed one hour before staining. Sections were rinsed 2 times in PBS for 5 minutes and for 1 minute in distilled water. Sections were put in pre-warmed (45 ˚C) cresyl violet solution for 20 minutes. Slides were then rinsed 2 times for 5 minutes in distilled water, followed by a 3 minute rinse in 90% ethanol and then 100% ethanol and cleared in xylene. Slides were mounted in DPX. Brightfield images were taken on a Zeiss Imager.D2 at 10 x magnification. Motor neurons were identified by morphology and size.

**Electron microscopy**

Sciatic nerves were isolated from mice and immediately fixed in 2.5% glutaraldehyde and 4% PFA for 2h at RT, and postfixed in 2.5% glutaraldehyde in PHEM buffer (1X, pH 7.4) overnight at 4°C. They were then rinsed in PHEM buffer and post-fixed in a 0.5% osmic acid for 2h at dark and room temperature. After two washs in PHEM buffer, the cells were dehydrated in a graded series of ethanol solutions (30-100%). The cells were embedded in EmBed 812 using an Automated Microwave Tissue Processor for Electronic Microscopy, Leica EM AMW. Thin sections (70 nm; Leica-Reichert Ultracut E) were collected at different levels of each block. These sections were counterstained with uranyl acetate 1.5% in 70% Ethanol and lead citrate and observed using a Tecnai F20 transmission electron microscope at 200KV in the CoMET MRI facilities, INM, Montpellier France.

Semithin (1 µm) cross section were stained with toluidine blue (Sigma-Aldrich, 89640-5G) and observed with a Nanozoomer Hamamatsu. The g-ratio was determined using the ImageJ GRatioCalculator plug-in (CIF, UNIL).

**Metabolomic**

Sciatic nerves of 4 months mice were dissected and immediately frozen in liquid nitrogen until metabolomics analysis. Samples were homogenized with 2 grinding cycles, each at 6600 rpm for 20 sec, spaced by 20 sec, using a Precellys homogenizer (Bertin Technologies, Montigny-le-Bretonneux, France) kept in a room at +4°C. The supernatant was recovered after centrifuging the homogenate and kept at -80°C until mass spectrometric analysis. Targeted metabolomic analysis was carried out using the AbsoluteIDQ kit p 180 (Biocrates Life Sciences AG, Innsbruck, Austria). This kit standardizes mass spectrometry quantification (Sciex QTRAP 5500 AB mass spectrometer, SCIEX, Villebon-sur-Yvette, France) of 188 metabolites including 40 acylcarnitines, 21 amino acids, 21 biogenic amines, 90 glycerophospholipids, 15 shingolipids and the sum of hexoses. The full list of individual metabolites is available at http://www.biocrates.com/products/research-products/absoluteidq-p180-kit. Flow injection analysis coupled with tandem mass spectrometry (FIA-MS/MS) was used for the analysis of carnitine, acylcarnitines, lipids and hexoses. Liquid chromatography (LC) was used for separating amino acids and biogenic amines before quantitation with mass spectrometry. Before statistical analysis, the raw metabolomics data were examined to exclude metabolites with more than 20% of concentration values below the lower limit of quantitation (LLOQ) or above the upper limit of quantitation (ULOQ). Before performing statistical analyses, data from each sample were normalized by the total ion current (TIC).

To determine if molecule content variations reflected differences in metabolism in between Wild-Type, control and mutant mice, we carried out a Principal Component Analysis (PCA) by using the R-package ade4 (*22*). This method is a classically used ordination method to summarize the patterns of variations among a large collection of samples, as it is well suited to contingency tables. As female's metabolism is more hormonal dependent than male, we excluded female mice to remove noise in the PCA. We thus performed the PCA on 15 male mice (5 WT, 5 controls and 5 mutants) with the full dataset of 161 molecules. Then we analyzed acylcarnityls profiles in control mutant and mutant mice using a paired two-tailed T-tests.

**Lactate assay**

To measure lactate content in sciatic nerves of Control and Mutant animals of 4 months and 12 months, equal-length nerves of 2 cm were dissected and immediately homogenized with Lysing Matix D tubes (MP Biomedical) in the lactate assay buffer and proceed according the instruction of the lactate assay kit (MAK064, Sigma-Aldrich).The absorbance at 570 nm was measured with microplate reader (CLARIOstar, BMG Labtech)

**Western Blot**

Sciatic nerves were dissected from mice, after removal of the epineurium and perineurium, the nerves were frozen in liquid nitrogen and conserved at -80°C. Protein were extracted from whole sciatic nerves after homogenization by sonication in standard RIPA lysis buffer and the protein concentration was measured using a BCA protein assay kit (Pierce). 20 µg proteins were directly analyzed by western blot using standard procedures with 12% SDS–PAGE and transferred on PVDF membranes for immunoblotting. Primary antibodies: Rabbit anti-Pkm1 1/2500 (NBP2-

14833, Novus Biologicals); Rabbit anti-Pkm2 1/2500 (SAB4200095, Sigma Aldrich); Mouse anti-α-β-Actin 1/10000 (C1.AC-15, #A1978, Sigma Aldrich). Secondary antibodies, Peroxidase Goat anti Rabbit or Mouse (H+L) (Jackson Immuno Research) were used at 1/10000 dilution.

**Semi-quantitative rtPCR**

Whole sciatic nerves from 2 and 10 days old mice (P2 and P10) or endoneuria from 1 (P28), 3 and 12 months-old mice were dissected at indicated time-points. Tissues were lysed using Trizol reagent and mechanical homogenization (TissueLyser II, Qiagen). Total RNA was extracted with the RNeasy lipid tissue kit (Qiagen), and RNA quality and concentration were verified by ND-1000 spectrophotometer (NanoDrop). cDNA was synthesized using 70-200ng of the RNA with the PrimeScript RT kit (Takara), following manufacturer's protocols. PKM1 and PKM2 splicing assays at different developmental time-points were conducted as previously described (*23*) using the following primers PKM-F: 5'-ATGCTGGAGAGCATGATCAAGAAGCCACGC-3' and PKM-R: 5'-CAACATCCATGGCCAAGTT-3' with PCR annealing step at 60°C during 35 cycles and using HotStart Taq Polymerase (Qiagen). Digestions of cleaned 502bp-PCR products (with DNA Extraction kit, Qiagen) by NcoI (NEB), producing approx. 250 bp bands revealing PKM1 transcript, and PstI (NEB), producing approx. 280+220 bp bands revealing PKM2 transcript), were performed for 2h at 37°C. Digested PCR products were resolved on a 2% agarose gel, imaged by ChemiDoc XRS+ system and quantified using Image lab software version 3.0 (Biorad). The ratio between detected intensity of uncut and cut products was used to assess relative amount of PKM1 and PKM2.

**Probes construction and AAV preparation**

pAAV-Laconic and pAAV-δGlu6 were obtained by digesting Laconic/pcDNA3.1(-) (gift from Luis Felipe Barros; Addgene plasmid #44238) (*15*) and pcDNA3.1 FLII12Pglu-700uδ6 (gift from Wolf Frommer; Addgene plasmid # 17866) (*17*) by BamH1/HindIII (NEB) and cloned into pAAV-MCS (Cell Biolabs, Inc.) under of a CMV promoter. Clones were validated by sequencing. pcDNA-mito-AT1.03 (from H. Imamura, Tokyo, Japan)(*18*) was digested with XhoI/HindIII (NEB), blunted and cloned into the CMV promoter controlled pAAV-MCS. Mitochondria-targeting tags were two tandem copies of CoxVIII. pAAV9 were produced at UPV, Universitat Autonoma de Barcelona or at INSERM U1089 University of Nantes, France.

***In vivo* injection in spinal cord**

The pups between 1 and 3 days were covered in aluminum foil and completely surrounded in ice for 3–4 min, until a completely cryoanesthetizia. Cryoanesthetized neonates were injected using a thin glass needle filled with colored viral and hold by a micromanipulator (IM—3C, Narishige Japan Group) directly in the spinal cord. 1 µL of the viral solution was injected slowly with short pressure pulses using a microinjector (Pneumatic Picopump PV820, World Precision Instruments) coupled to a 3 MHz function pulse generator (GFG8215, Langlois). The injection site was then cleaned with betadine (Vetoquinol, cat. No. 3042413) and pups were warmed in hands. When fully awake, pups were put back to the mother and the littermates.

**Imaging and stimulation of saphenous nerves in living mice**

3-4 weeks after the viral injection, mice were anesthetized with a constant low (1.5 l/min) of oxygen+ 5% of isoflurane in an anesthesia induction box (World precision Instrument, Ref.EZ-B800) for 5 min. Thereafter the anesthesia was maintained with a mask delivering 2% isoflurane at 0.8 L/min. The eyes were protected with Ocry-gel (TVM, cat. No. 48026T613/3). The mouse was placed on the back in a silicone mold, the hind leg was shaved and the paws were immobilized using small pins. The incision area was disinfected with betadine (Vetoquinol, cat. No. 3042413). After incision of the skin of the thigh, connective tissue was carefully removed to expose the saphenous nerve. For electrophysiological experiments, the saphenous nerve was lifted up in the middle of the thigh and isolated electrically from the surrounding muscle tissue using a plastic trip. A pair of stimulating platinum electrodes (World precision Instrument, PTM23B05KT) held by micromanipulators (U-31CF, Narishige) was carefully placed under the saphenous nerve, at the ankle. The recording electrode (AD Instruments, MLA 1203) was placed at the groin. A reference needle electrode was inserted in the groin area and a ground electrode is placed in the tail. Once the electrodes were positioned the animal was transferred in a chamber at 37°C under the two-photon microscope (LSM 7 MP OPO, Zeiss), electrodes are connected to a Powerlab 26T (AD Instrument; ML4856). Supramaximal stimulus (between 4 and 12 mA, 150 µs) was delivered at a frequency of 10 Hz during 5 min (*24*). Time-lapse of the middle portion of the exposed nerve in the thigh were acquired every 5 min during 1 h with 20X objective lens (LD CApochromat, 421887-9970, Zeiss. For Laconic, δGlu6, and mito-ATeam alike, a 870 nm excitation wavelength was used to obtain both emission wavelength 475 nm for mTFP (Laconic) or CFP (δGlu6, mito-Ateam) and emission wavelength 527 nm for Venus (Laconic, mito-ATeam) or YFP (δGlu6).

**Image analysis**

We used ImageJ software to analyze the relative lactate, glucose or mitochondrial ATP levels in peripheral axons. The acquired images for each wavelength (em. 475 nm for mTFP or CFP and em. 527 nm for Venus or YFP) were aligned using the Template Matching plugin. We defined a Region of Interest (ROI) encompassing the cytosolic area of one axon or encompassing all labelled mitochondria of one axon and the mean fluorescent intensity in the ROI was measured on both images. These light intensities were then corrected for background light intensity determined as an area within the nerve where no fluorescent signal from the viral probe can be observed. The Citrine/CFP (δGlu6) or Venus/mTFP (Laconic) or Venus/CFP (mito-Ateam) ratio was then calculated from the 2 values for mean light intensity.

The MTrackJ plugin in ImageJ was used for analysis of mitochondrial movement. For at least 5 images over a time span of 20 minutes, the location of the same mitochondrion was marked, the distance between marked locations was measured and migration velocity was calculated. n= 65 (Control before stimulation), 75 (Mutant before stimulation), 76 (Control after stimulation), 45 (Mutant after stimulation) mitochondria.

**Ex vivo nerve electrophysiology**

Sciatic nerves were dissected out and transferred into oxygenated artificial cerebrospinal fluid (ACSF) containing 126 mM NaCl, 3 mM KCl, 2 mM $CaCl_2$, 2 mM $MgSO_4$, 1.25 mM $NaH_2PO_4$, 26 mM $NaHCO_3$, and 10 mM dextrose, pH 7.4-7.5. The nerves were desheated, and cut into 2 cm segments. Nerves were then placed in a three compartment recording chamber and perfused (1-2 ml/min) in 36°C ACSF equilibrated with 95% $O_2$-5% $CO_2$. The distal end was stimulated supramaximally (40 µs duration) through two electrodes isolated with Vaseline, and recordings were performed at the proximal end. Signals were amplified and digitized at 500 kHz. Measurements were made once the effects had reached a steady state. The delay and duration of compound action potentials (CAPs) were calculated at half the maximal amplitude and at the maximal amplitude. For recruitment analysis, the amplitude of CAPs was measured and plotted as a function of the stimulation intensity. For refractory period analysis, two stimuli were applied at different intervals, and the amplitude of the second CAP was measured and plotted as a function of the stimulus interval. To ensure that the amplitude of the second response was accurately assessed, the first response was subtracted from all the recordings. For train stimulation, nerves were stimulated for 200ms at frequencies ranging from 100 to 1 kHz. The amplitude of each evoked CAP was measured and plotted as a function of time. Conduction velocities were estimated from latencies.

**Statistical significance**

*= P value<0.05; **= P value<0.01; ***= P value<0.001.

**Supplementary Figures and Table:**

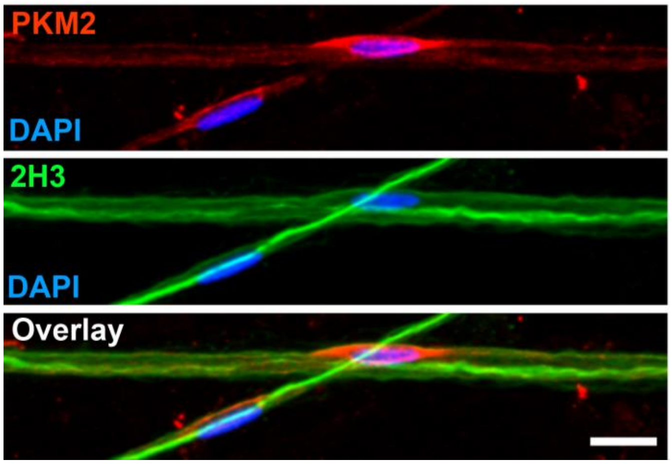


**FIGURE S1 PKM2 is enriched in the perinuclear cytoplasm of mSC.** Immunostaining of mature mSC in teased mouse sciatic nerve (1 month old) fibres for PKM2, axonal 2H3 and with nuclear DAPI show the enrichment of PKM2 in the perinuclear cytoplasm of mSC.
Scale bar= 10μm.

**FIGURE S2**

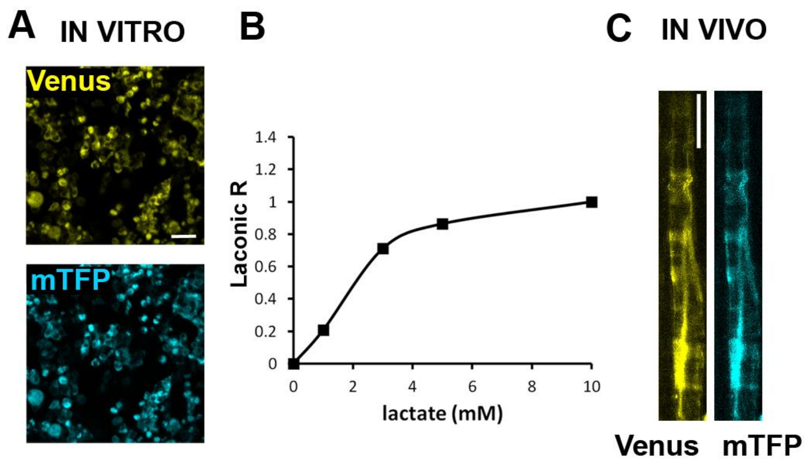


**FIGURE S2 PKM2 is enriched in the perinuclear cytoplasm of mSC.**
Immunostaining of mature mSC in teased mouse sciatic nerve (1 month old) fibres for PKM2, axonal 2H3 and with nuclear DAPI show the enrichment of PKM2 in the perinuclear cytoplasm of mSC.
Scale bar= 10μm.

**FIGURE S3**

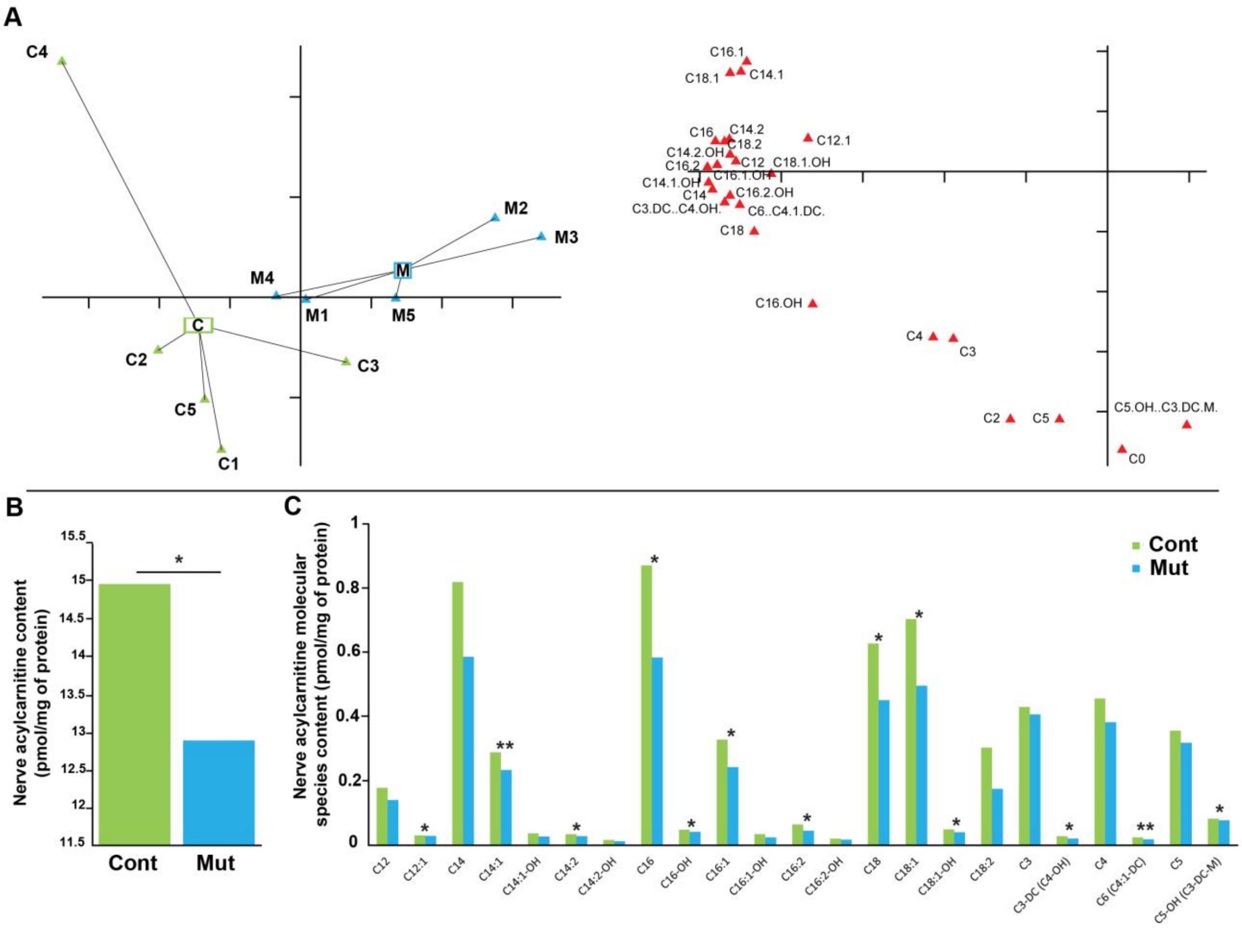


**FIGURE S3 Metabolomic analysis of Control and Mutant sciatic nerves. A-** A PCA performed on acylcarnithines separates Mutant (M) from Control (C) mouse nerves with strong eigenvalues (horizontal axis 64% and vertical axis 17%). Mutant and Control are mainly separated along the axis-1 showing that Control are associated with stronger occurrences of long chain acylcarnithines. n=5 male mice per genotype. **B-** Nerves of Mutant mice contain significantly less acylcarnitines than Controls. **C-** The systematic analysis of acylcarnitines shows a significant decrease of long-chain molecular species in Mutant vs Control nerves.

n=5 male mice per genotype. Female metabolism was noisier due to hormonal variability, so we excluded female mice from the analysis. Statistical test show paired two-tailed Student T-test. * = P<0.05.

### FIGURE S4

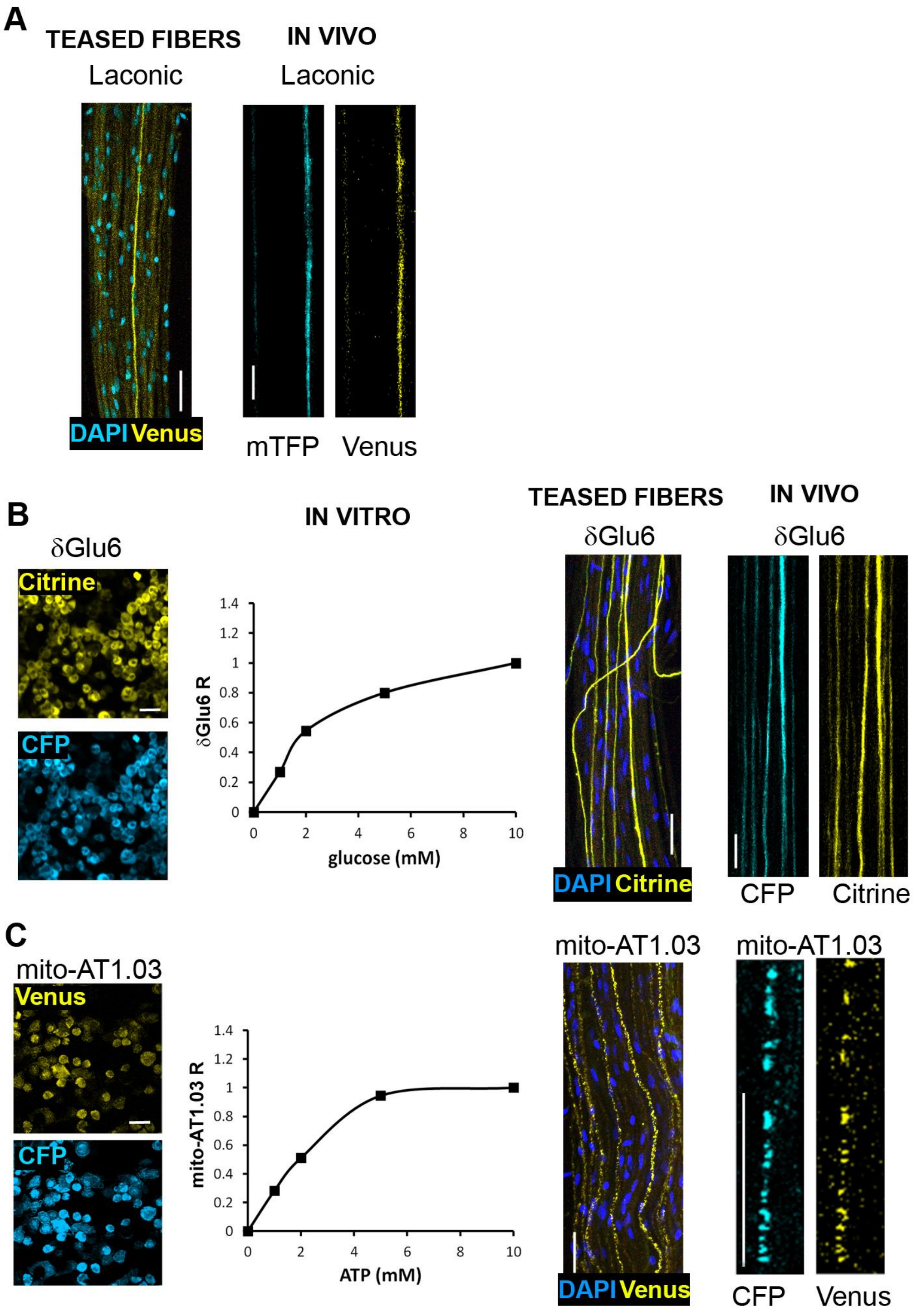


**FIGURE S4 Functional validation of the fluorescent probes. A- left:** HEK293 cells were transfected with pAAV-Laconic and imaged for Venus (yellow) and mTFP (blue)(scale bar= 50 µm). Then Venus/ mTFP fluorescence ratio (Laconic R) was measured at different concentration of lactate (0, 1, 3, 5, 10 mM) showing a correlation between increasing amount of lactate and Laconic fluorescence ratio (Graph). **Right:** AAV9 expressing Laconic under a CAG promoter was injected in the spinal cord of newborn mice and one month later Mutant and control animals were treated with Tamoxifen. Three to five weeks later, sciatic nerve fibers were teased on a glass slide. Laconic (Venus, yellow) is expressed in long never-ending axons (scale bar= 100μm). DAPI (blue) shows nuclei of the surrounding cells. In addition, the saphenous nerve of anesthetized mice expressing Laconic was exposed under the lens of a multi-photon microscope and both Venus (yellow) and mTFP (blue) fluorescence were recorded in order to measure Laconic fluorescence ratio *in vivo* (scale bar= 10μm). **B**- Similar experiments were done as in **A** for δGlu6 fluorescent probe. δGlu6 fluorescence ratio (Citrine/CFP) is proportional to glucose concentration (0, 1, 2, 5, 10mM). The probe is expressed in axons crossing the sciatic nerve and the exposed saphenous nerve allows to record this ratio in axons of living anesthetized mice. **C**- Similar experiments were done as in **A** for mito-Ateam (mito-AT1.03). Mito-Ateam fluorescence ratio (Venus/CFP) is proportional to ATP concentration (0, 1, 2, 5, 10mM). The probe is expressed in mitochondria (dotted pattern) of axons crossing the sciatic nerve and the exposed saphenous nerve allows to record this ratio in axonal mitochondria of living anesthetized mice.

#### FIGURE S5

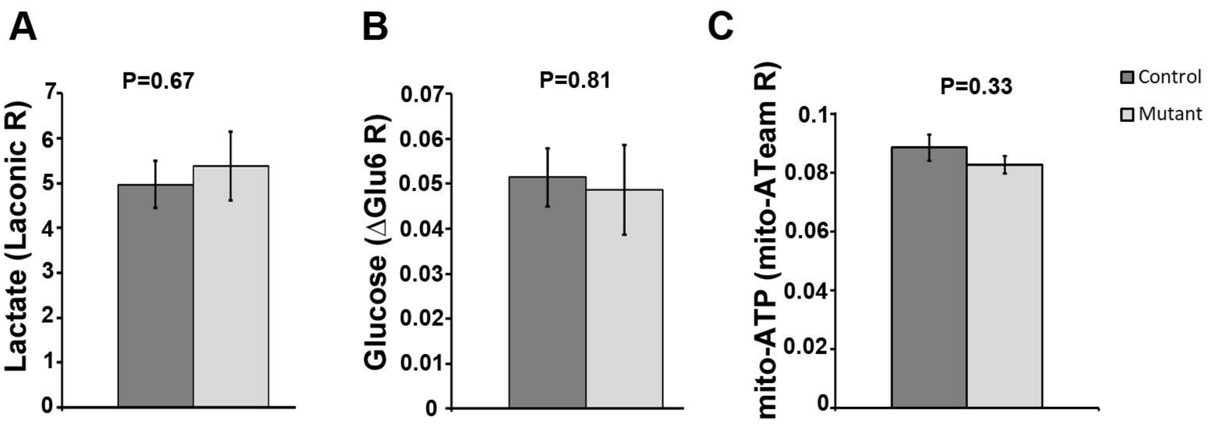


**FIGURE S5 Lactate, glucose and mitochondrial ATP levels in resting axons of Mutant and Control saphenous nerves are not different.** As described in Figure S3, Laconic (**A**), δGlu6 (**B**) and mito-Ateam (**C**) fluorescence ratios were measured in resting axons of the saphenous nerves of Mutant and Control mice. No significant differences could be detected in these conditions. P-values show two-tailed unpaired Student T-test. A: n=15 axons in 7 Control mice, 7 axons in 4 Mutant mice. B: n=9 axons in 3 Control mice, 6 axons in 3 Mutant mice. C: n=14 axons in 5 Mutant mice, 16 axons in 5 Control mice. Error bars represent SEM.

FIGURE S6

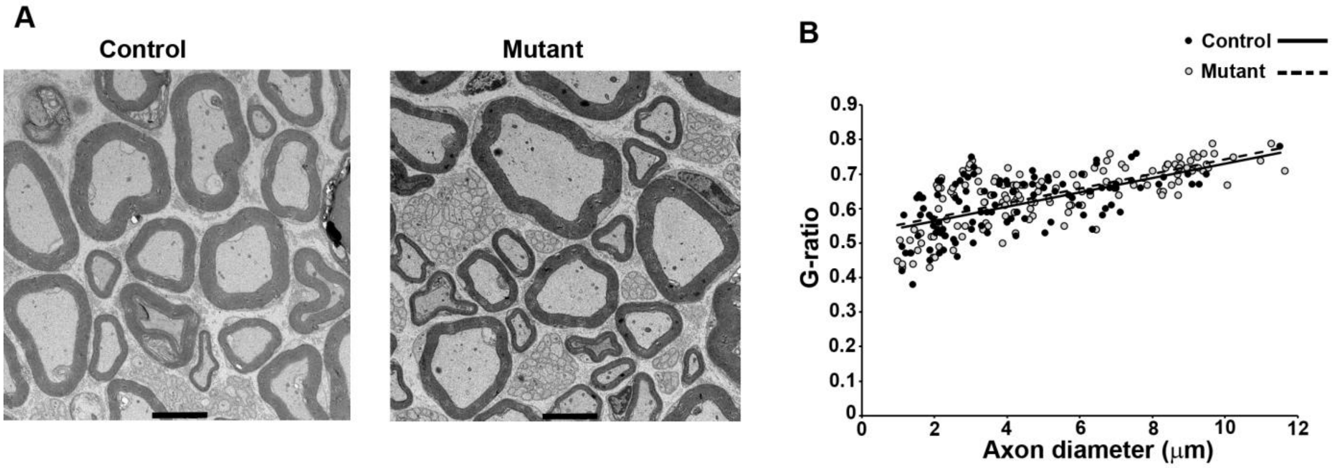


**FIGURE S6 Electron microscopy analysis of Mutant and Control mouse sciatic nerves. A-** Ultrathin sections of Control and Mutant mouse sciatic nerves (12 months old) did not reveal any structural difference between genotypes. Scale bars= 4μm. **B**- Using semi-thin section, the ratio axon diameter on full fiber diameter (G-ratio) was measured and plotted relative to the axon diameter. No difference was seen for the regression lines between genotypes showing that myelin diameter is no affected by the deletion of PKM2 in mSC. n= 3 animals for both genotype.

#### FIGURE S7

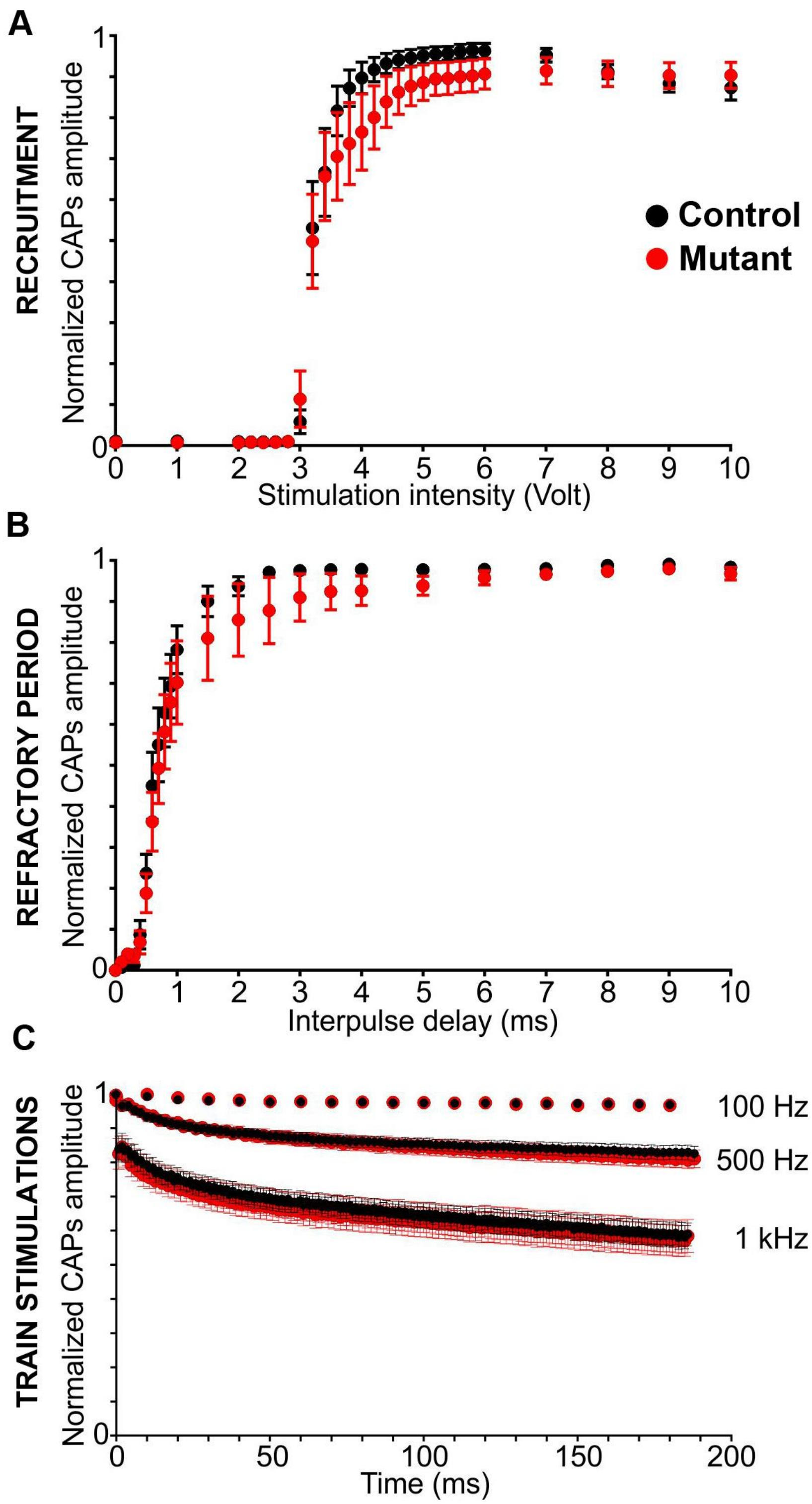


**FIGURE S7**

**Electrophysiological analysis of Control and Mutant sciatic nerves show no significant alteration of the action potential propagation along axons.** **A, B**- The recruitment (**A**) and refractory period (**B**) of sciatic nerves from Mutant mice (n = 10 nerves from 5 mice) is not significantly different from those of Control mice (n = 10 nerves from 5 mice) (P>0.05 by two-tailed t-tests for two samples of equal variance). **C**- Nerves were stimulated with train of stimuli ranging from 100 to 1 kHz in order to monitor the sustainability of the response. No difference was observed between Control and Mutant mice. Error bars represent SEM.

**TABLE S Electrophysiological characteristics of Control and Mutant mice.**

| | Control | Mutant |
|---|---|---|
| Amplitude (mV) | 2.5 ± 1.8 | 3.2 ± 2.5 |
| Duration (ms) | 0.34 ± 0.06 | 0.37 ± 0.09 |
| $CV_{V½}$ (m.s$^{-1}$) | 52.1 ± 15.8 | 45.2 ± 16.1 |
| $CV_{Vmax}$ (m.s$^{-1}$) | 33.8 ± 8.5 | 30.6 ± 9.4 |
| n | 10 (5 animals) | 10 (5 animals) |

#### FIGURE S8

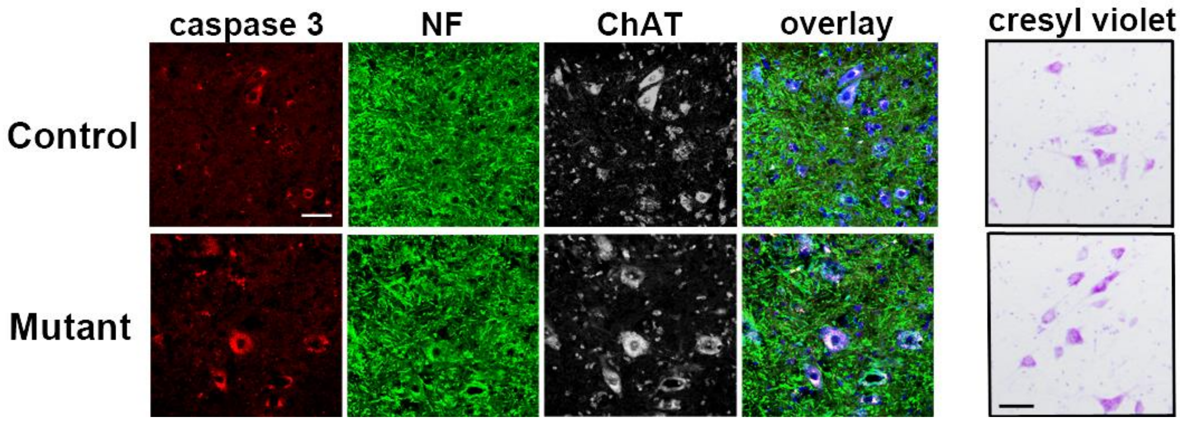


**FIGURE S8 Mutant mouse motor neurons display neuronal stress marker Caspase 3.** Spinal cord cryosections of Mutant and Control mice were stained with cresyl violet or immunostained for neuronal stress marker Caspase 3 (red), neuronal marker Neurofilament (NF, green) and motor neurons marker ChAT (blue). While Neurofilament staining is not different, more motor neurons express Caspase 3 in Mutant mice than in Control mice. Scale bars= 50µm.
